# Transcription templated assembly of the nucleolus in the *C. elegans* embryo

**DOI:** 10.1101/2024.06.06.597440

**Authors:** Nishant Kodan, Rabeya Hussaini, Stephanie C. Weber, Jane Kondev, Lishibanya Mohapatra

**Affiliations:** Department of Physics, New York University, New York, NY 10003, USA; School of Physics and Astronomy, College of Science, Rochester Institute of Technology, Rochester, NY 14623, USA; Department of Biology, McGill University, Montreal, QC H3A 1B1, Canada; Department of Physics, McGill University, Montreal, QC H3A 2T8, Canada; Department of Physics, Brandeis University, Waltham, MA 02454, USA

**Keywords:** Nucleolar assembly, transcription-templated assembly, membraneless organelles

## Abstract

The nucleolus is a multicomponent structure made of RNA and proteins that serves as the site of ribosome biogenesis within the nucleus. It has been extensively studied as a prototype of a biomolecular condensate whose assembly is driven by phase separation. While the steady-state size of the nucleolus is quantitatively accounted for by the thermodynamics of phase separation, we show that experimental measurements of the assembly dynamics are inconsistent with a simple model of a phase-separating system relaxing to its equilibrium state. Instead, we show that the dynamics are well described by a model in which the transcription of ribosomal RNA actively drives nucleolar assembly. We find that our model of active transcription-templated assembly quantitatively accounts for the rapid kinetics observed in early embryos at different developmental stages, and for different RNAi perturbations of embryo size. Our model predicts a scaling of the time to assembly with the volume of the nucleus to the one-third power, which is confirmed by experimental data. Our study highlights the role of active processes such as transcription in controlling the placement and timing of assembly of membraneless organelles.

**Significance statement:** How membraneless organelles like nucleolus assemble within cells is not well understood. Recent experiments suggest that transcription of ribosomal RNA actively drives nucleolar assembly. Our proposed model of active transcription-templated assembly quantitatively accounts for the rapid kinetics observed in early worm embryos at different developmental stages. Further, it predicts a scaling of the time to assembly with the volume of the nucleus that is confirmed by experimental data. This work describes how active processes such as transcription can control the placement and timing of assembly of membraneless organelles.

## Introduction

The contents of a cell are compartmentalized into organelles with specific functions. Traditionally organelles have been defined as compartments surrounded by a membrane, like the nucleus, mitochondria and cytoskeletal structures like cilia, but more recently membraneless compartments have been described, like P granules (1), nucleoli (2), and stress granules (3). Despite the absence of a membrane, these organelles are able to maintain a coherent structure while their components rapidly exchange with the cytoplasmic pool. Phase separation has been identified as the key physical process that maintains the distinct composition of membraneless organelles(4). While phase separation provides the thermodynamic driving force that stabilizes these cellular structures against breakdown by diffusion of their molecular components, how their assembly is regulated in space and over time, remains an open question.

The nucleolus has emerged as a model system for studying membraneless organelles(5). It is composed of both RNA and protein components and serves as the site of ribosome biogenesis within the cell nucleus. In early *C. elegans* embryos, nucleolar proteins are rapidly imported into the nucleus, where they condense to form dozens of small extranucleolar droplets (ENDs) (6). Over time, ENDs coarsen into two large nucleoli, which are located at the two ribosomal DNA sites present in diploid cells. Nucleoli ultimately dissolve as cells enter mitosis, and the soluble nucleolar components are partitioned proportionally to daughter cells. As the cells become smaller with successive divisions, the nuclei scale proportionally and so does the maximal size of both nucleoli (7).

*C. elegans* embryogenesis occurs in an egg of fixed size, such that blastomeres become progressively smaller through reductive divisions. The pattern of divisions is invariant, giving rise to a well-defined cell lineage (8). In wildtype embryos, nucleoli do not appear consistently until the 8-cell stage (7), which consists of 4 anterior cells of the “AB” lineage (ABar, ABal, ABpr, ABpl); the ventral founder cells MS and E; and the posterior cells C and P3. In the subsequent 16-cell stage, there are 8 AB cells, and two cells in each of the MS and E lineages (MSa, MSp, Ea, and Ep). Cells in the AB lineage divide quickly (cell cycle ∼15 min), while cells in the MS and especially E lineages divide more slowly (cell cycle ∼25 min). This natural variation in cell cycle time allows us to compare nucleoli that assemble and disassemble rapidly (AB lineage) with those that persist at an apparent steady-state for an extended period of time (E lineage). We track the dynamics of nucleolar assembly in embryos expressing a fluorescent fusion of the conserved nucleolar component fibrillarin (FIB-1::GFP) as a function of developmental stage and in the presence of different RNAi-mediated perturbations of embryo (and nucleolar) size (7).

Using fluorescence microscopy, a previous study reported that the total amount of nucleolar material (represented by FIB-1::GFP protein) in an embryo is equal to what is maternally loaded from the oocyte until the 128-cell stage, thus suggesting that there is no significant zygotic contribution of FIB-1 protein in these early embryos (7). Using experiments and theory, they showed that maximum nucleolar size scales with nuclear size during early embryonic divisions and proposed a “limiting-pool mechanism” to explain this size scaling phenomenon. This mechanism is based on a simple idea that an organelle grows until the pool of its building blocks is depleted to the point where the rates of assembly and disassembly of the organelle are balanced (9). This mechanism also makes several predictions for the kinetics of assembly(10), that we analyze quantitatively here.

In addition to a limiting pool of protein components, RNA and/or the process of transcription may also contribute to nucleolar assembly. Berry et al found that at the 8-cell stage, within the AB lineage, nucleolar proteins first condense into ENDs, which dissolve and coalesce to supply material for the assembly of the two nucleoli. They also report a co-localization of nascent rRNAs with the nucleolus at the 8-cell stage using fluorescence *in situ* hybridization (FISH) (6) and note that nucleoli do not form when transcription is inhibited. This striking observation prompted the authors to propose a theoretical model describing nuclear bodies as liquid phase droplets. By considering the nucleus as a fluid consisting of nucleolar proteins in the nucleoplasm which coalesce into droplets within a region defined by the transcribed rRNA, they showed that the nucleolus forms if protein-protein interactions considered within this region are stronger than in the bulk. While this paper elegantly showed a qualitative agreement with the experimental data in terms of the differential kinetics of the assembly of ENDs and nucleoli, the quantitative relationship between the dynamics of transcription and nucleolar assembly remains unexplored.

Here, we explore how the process of RNA synthesis contributes to the kinetics of membraneless organelle assembly in cells.

We find that theoretical predictions of the limiting pool mechanism cannot explain quantitatively the dynamics of nucleolar assembly observed in experiments. Instead, we consider models of transcription-driven assembly of the nucleolus in which RNA serves as a nucleator of the condensed phase (11–13). We estimate model parameters from experimental data and use them to generate growth trajectories of two nucleoli assembling in a limited pool of building blocks. By quantitatively comparing our simulated trajectories with experimental data from different cell stages and lineages, we find that our active transcription-templated growth model is able to quantitatively account for nucleolus assembly in WT and in different RNAi conditions. Further, we propose experiments to test our model predictions and quantitatively study the role of transcription in the assembly of the nucleolus. Overall, our results suggest a mechanism by which cells can use the production of nucleators to control organelle assembly in space and time.

### Passive versus active transcription-templated assembly of the nucleolus

In early *C. elegans* embryos, in the process of cell division, nucleolar proteins are passed from the mother cell to the two daughter cells. Nucleoli form in each daughter cell using the inherited material and then disassemble as the cell prepares for the next division. As the developmental stages progress, and the cells get smaller, nucleolar material continues to be divided proportionally among the daughter cells. Consequently, the nucleoli in each cell get smaller in size. Here we analyze two simple models of the assembly process with the goal of describing the time evolution of nucleolar size. We build our models by mathematically describing three key experimental observations (6, 7): one, that nucleolar assembly is nucleated by the ribosomal RNA (rRNA), which is transcribed from ribosomal genes; two, that the growth of the nucleolus is controlled by a limiting pool of one of its components; and, three, that the condensation of nucleolar material into small droplets (ENDs), which is seen in the absence of rRNA, proceeds at a constant rate.

The ultimate goal of our analysis is to understand the role of rRNA synthesis in the assembly of the nucleolus by comparing predictions from mathematical models to quantitative measurements of nucleolar size with time. We consider two models of rRNA driven nucleolar assembly: one in which the nucleolus assembles on a pre-formed rRNA template (the “passive” model), and another where the rRNA template is synthesized while the nucleolus is simultaneously being assembled (the “active” model). In the active model, the process of transcription of rRNA effectively drives the assembly of the nucleolus, with the nascent rRNA serving as nucleators on which the ENDs condense. In this section, we describe both models and compare their predictions for nucleolar assembly with data obtained in experiments.

In the absence of ribosomal RNA, nucleolar FIB-1 proteins condense into droplets whose size grows linearly with time (6). We mathematically describe this observation using the rate equation

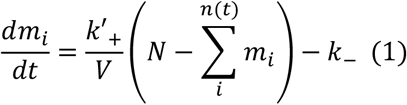

where *m*_*i*_ is the size of individual droplets, measured in the number of nucleolar subunits (GFP labels FIB-1 molecules in the experiment) and *n*(*t*) is the number of droplets at time *t*. The time-dependent rate constants, 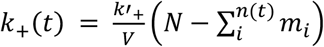 is the rates at which FIB-1 molecules are incorporated, where the second order rate constant *k*′_+_ describes the rate of assembly of a droplet from the free pool of nucleolar subunits and the concentration of free subunits is 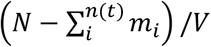, where *N* is the total number of nucleolar subunits within the nucleus of volume *V*. The first-order disassembly rate constant, *k*_−_, describes the process of subunits detaching from the droplets and rejoining the free pool. To capture the experimentally observed linear-in-time growth of the nucleolar droplets in the absence of rRNA, we assume that both rate constants do not change with the size of the nucleolar droplets nor with time.

To account for the experimental observation that the nucleolus assembles at the site of ribosomal RNA transcription, we assume that the rRNA provides nucleation sites for the growth of the nucleolus. At each nucleation site, a droplet forms following Equation 1 and the size of the nucleolus *M* is simply the sum of the mass of all the droplets, *M* = ∑_*i*_ *m*_*i*_.

The equation to describe the growth of nucleolus *M*(*t*) is obtained by summing Equation 1 over all *n*(*t*) nucleolar droplets:

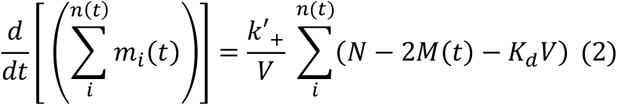

where 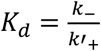 is the dissociation constant of the condensation reaction in which the droplet grows by adding one subunit from the nucleoplasm. The concentration of free nucleolar subunit pool is 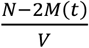 since there are two nucleoli per cell in worm embryos, each being size M(t).

We can now consider two situations, one, where the nascent rRNA amount reaches its steady state well before the nucleolus assembles. In this case the number of nucleolar droplets *n*(*t*) = *n*_*max*_ is independent of time, and we refer to this as the Passive model. The other possibility is that rRNA transcription unfolds concomitantly with nucleolar assembly, in which case the number of droplets is time-dependent and proportional to the amount of nascent rRNA. This is the Active model of nucleolus assembly.

*Passive transcription-templated assembly:* In this model we assume, that the amount of nascent rRNA is fixed during the process of nucleolar assembly. This is to be expected if the first RNA polymerase completes transcription before the nucleolus reaches the minimum size that we can experimentally detect. At the point in time when we detect the growth of the nucleolus we expect in this case that the rRNA gene is saturated with RNA polymerases. The number of nascent rRNAs is at its maximum, *n*(*t*) = *n*_*max*_

The size of a nucleolus is the sum of the sizes of all the nucleolar droplets, 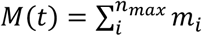. Using this in Equation 2, we arrive at:

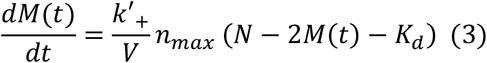

for the rate equation that describes the growth of a single nucleolus.

The solution of Equation 3 is 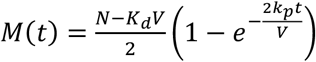, where *k*_*p*_ = *k*′_+_ *n*_*max*_; it predicts that at early times, when 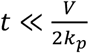, *M*(*t*) grows linearly with respect to *t*, as observed experimentally for the ENDs that form in the absence of rRNA. Furthermore, at long times, *t* ≫ *V*/2*k*_*p*_ Equation 3 predicts a steady state size of the nucleolus,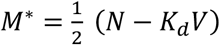. Furthermore, using *N* = *CV*, where *C* is the total concentration of subunits in the nucleus, we can write the expression for the total size of nucleoli at steady state as 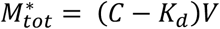, which is just twice the size of one nucleolus. This linear relationship between total size of the two nucleoli and volume of the nucleus (*V*) was previously put to a quantitative test (7). Later in this paper, we use this result and the solution of Equation 3 for *M*(*t*) to extract the parameters (*N, K*_*d*_ and *k*_*p*_) for the passive model using experimental data. It is important to note that the model described by Equation 3 is a much-simplified version of the model of nucleolar growth described previously (6) which made use of the free energy functional of the condensation explicitly and its dependence on the position-dependent subunit concentration within the nucleus. Still, both the published model and the one we adopt rest on the same biophysical assumptions described above.

*Active transcription-templated assembly:* In early worm embryos, the time scales of transcription (see supplementary materials) and the formation of nucleolus are similar (∼ 10 min), making it likely that the first RNA polymerase to complete transcription of rRNA is still transcribing as the nucleolus is forming.

Using 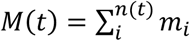 and assuming that there are *n*(*t*) independent nucleolar droplets that make up the nucleolus at time *t*, from Equation 2, we get

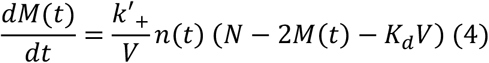

Therefore, the only difference between this active model (Equation 4) and the previously described passive model (Equation 3) is that in the passive case *n*(*t*) is assumed to be time-independent.

In order to compute the amount of nascent rRNA, we model transcription of rRNA as a two-step process (Figure 1A): (1) initiation – in which the RNA polymerase is recruited to promoter DNA and initiates transcription at rate *α*_*I*_, and (2) elongation – in which synthesis of nascent rRNA by the RNA polymerase proceeds at rate *α*_*E*_, as the polymerase moves along the rRNA gene. Inspired by experiments on the dynamics of transcription in the early fly embryo (14), we assume that both processes are described by a constant rate leading to a linear in time rise in the number of nascent transcripts, and a quadratic in time growth of the total amount of rRNA (measured in number of transcribed nucleotides), i.e. *n*(*t*) = *α*_*I*_*α*_*E*_*t*^2^/2. Note that in the active model, we are in a regime where we assume that the rRNA gene is saturated with polymerases, and the number of nascent rRNAs is no longer time-dependent, reaching a fixed value, i.e., *n*(*t*) = *n*_*max*_.

**Figure 1:**
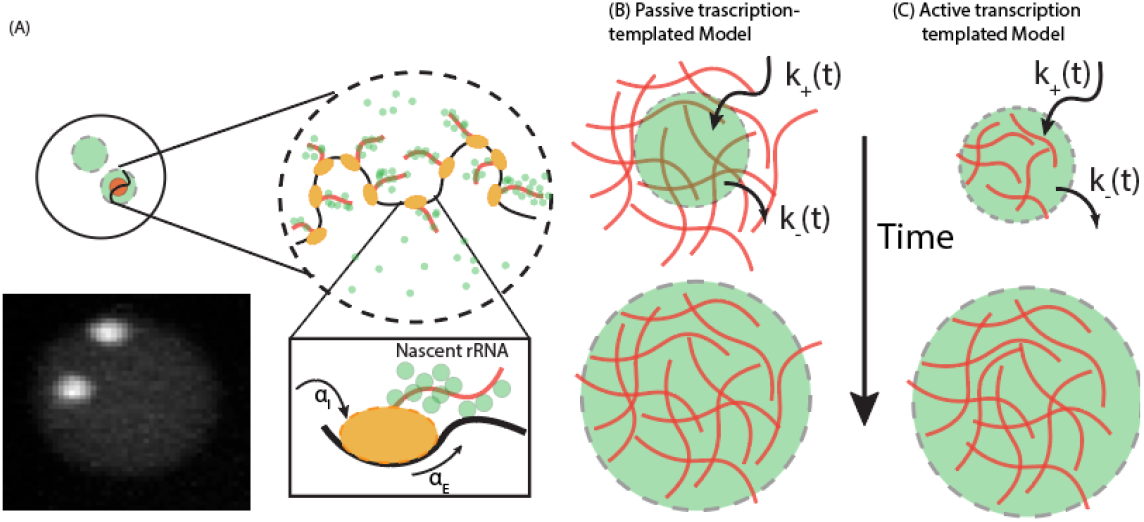
Models of transcription-templated nucleolar assembly. (A) Nucleoli (white regions in the fluorescent image) assemble at sites of rRNA transcription. Nascent rRNA molecules (red), produced during transcription, provide nucleation sites for condensation of FIB-1 molecules (green). Transcription of rRNA is initiated with rate *α*_*I*_ and the nascent transcript is elongated at rate *α*_*E*_. (B) Passive transcription-templated model: nucleoli assemble at pre-formed rRNA. (C) Active transcription-templated model: nucleoli assemble while rRNA is being transcribed. The time-dependent rate constants, *k*_+_(*t*) and *k*_−_( *t*), are the rates at which FIB-1 molecules are incorporated and leave the growing nucleolus.

Hence the change in the size of the nucleolus (given by the number of nucleolar subunits) in unit time is

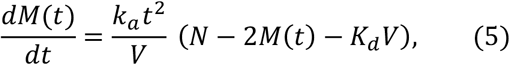

where we define, 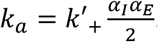. At steady state, (*N* − 2*M*^*^− *K*_*d*_*V*) = 0 and hence,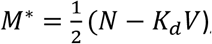, which is identical to the prediction from the passive model, whereas the time-dependent solution of Equation 5 is 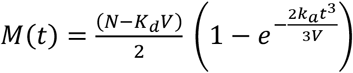.

Notably, while the two models predict the same nucleolar size at steady state, they differ qualitatively in their predictions for the time dependence of the size of a growing nucleolus (Figure 2B). The passive model predicts a linear dependence of the time to assembly on the volume of the nucleolus, while the active model predicts this time to scale with volume to the one-third power. In the next section, we use experimental data on the dynamics of nucleolar growth to discern between these two models, and find that the active model does a much better job of accounting for the measurements.

**Figure 2:**
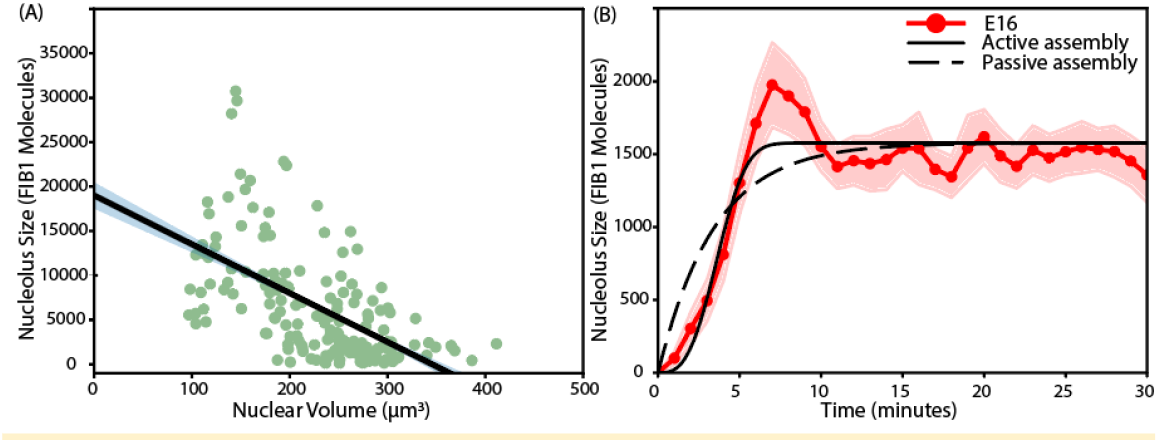
Nucleolar size and assembly dynamics: Theory-experiment comparison. (A) Comparison between the measured steady-state size of the nucleolus and the volume of the nucleus within which it assembled. Data points (green circles) correspond to AB8 nuclei from different RNAi conditions. The theory curve (black) is the prediction of both models of nucleolar assembly at steady state. (B) Dynamics of nucleolar assembly in E16 (red dots and shaded curve) compared to the predictions of the passive (dashed black line) and active model (solid black line) of rRNA templated assembly. The experimental data show the mean (red dots) and standard error of the mean (shaded red) over 8 independent measurements.

### Nucleolar assembly in the *C. elegans* embryo is actively driven by rRNA transcription

To compare predictions for nucleolar assembly from the two models, we first fit model parameters associated with the steady state from AB lineage cells: 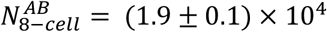 molecules, and 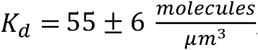, using data reported in ref (7) (Figure 2A and SI). We assume that the parameter *K*_*d*_ remains the same at the 8 and 16-cell stages, and that the different lineages at the 8-cell stage all have the same concentration of subunits. This allows us to compute the total number of molecules *N* in each cell type, as shown in Table 1.

**Table 1:**
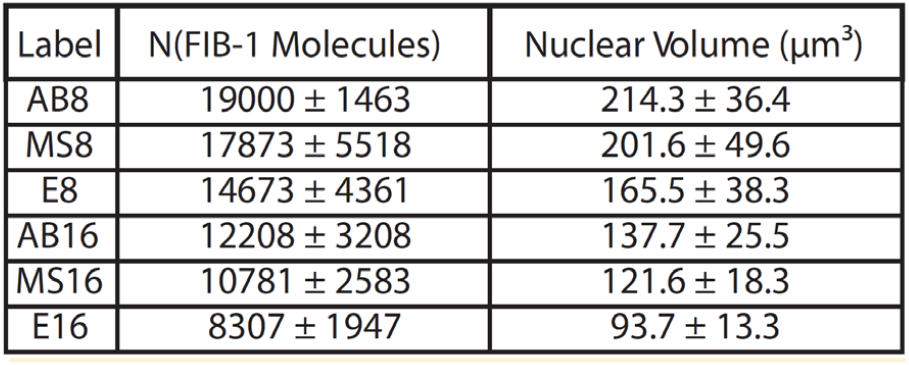
Predicted number of FIB-1 molecules (N) and measured nuclear volume (with standard deviation) for each lineage at the 8 and 16-cell stages. The AB8 value is obtained by fitting the steady state prediction of the models to the data (Fig 2A), and all other values of N are predictions from the model.

Next, we focus on the E lineage at the 16-cell stage due to its extended cell cycle time. Unlike earlier stages in which nucleoli quickly dissolve after reaching a maximum size, 16-cell E-lineage nucleoli persist around the same size for ∼ 20 min before dissolving, providing an apparent st eady-state (Figure 2B). Using *N* and *V* for E16 cells (i.e. Ea and Ep), we fit predictions for *M*(*t*) from the two models with measured fluorescence data and obtain fitting parameters *k*_*p*_ = 13.6 ± 1.8 *μm*^3^/*min* for the passive model and 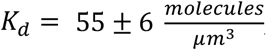 for the active model (see SI). Comparing the predicted curves, we note that the two models show different time dependence while leading to the same steady state, as expected. Notably, the active model is able to capture the concavity seen at early time (≤ 5 min), suggesting that rRNA template accumulates while the nucleolus is assembling. We next use the active model to predict and compare nucleolar assembly in different stages of development and following different RNAi perturbations.

### Active model of nucleolar assembly is in quantitative agreement with experiments at different stages of development and for different RNAi perturbations of the *C. elegans* embryo

At the 8-cell stage, there are 4 AB, 1 MS and 1 E cells which double in the 16-cell stage (Figure 3A). (The C, D, and P lineages do not consistently assemble nucleoli at these stages and were not included in this analysis.) In the previous section, we used E16 cells to extract model parameter *k*_*a*_ for active transcription-templated assembly. Since *k*_*a*_ depends on transcription rate parameters, we assume that the parameter *k*_*a*_ is the same for all cell stages and lineages examined. By using the number of molecules and nuclear volumes for each lineage (See table 1), we compare the theoretical predictions for the growth of the nucleolus to data obtained from fluorescence microscopy experiments (see Methods). Remarkably, we find that our predictions of active transcription-templated assembly capture the early stages of growth (<=5 min) in the nucleoli at both cell stages for all lineages examined (figures 3 and S3). In making this comparison, we introduced an additional parameter, namely the delay time *t*_0_ after the beginning of the developmental stage, at which the transcription of the rRNA genes and therefore the growth of the nucleolus starts (see Methods). Notably, all the other parameters appearing in Equation 3 are at this point of the analysis fixed.

**Figure 3:**
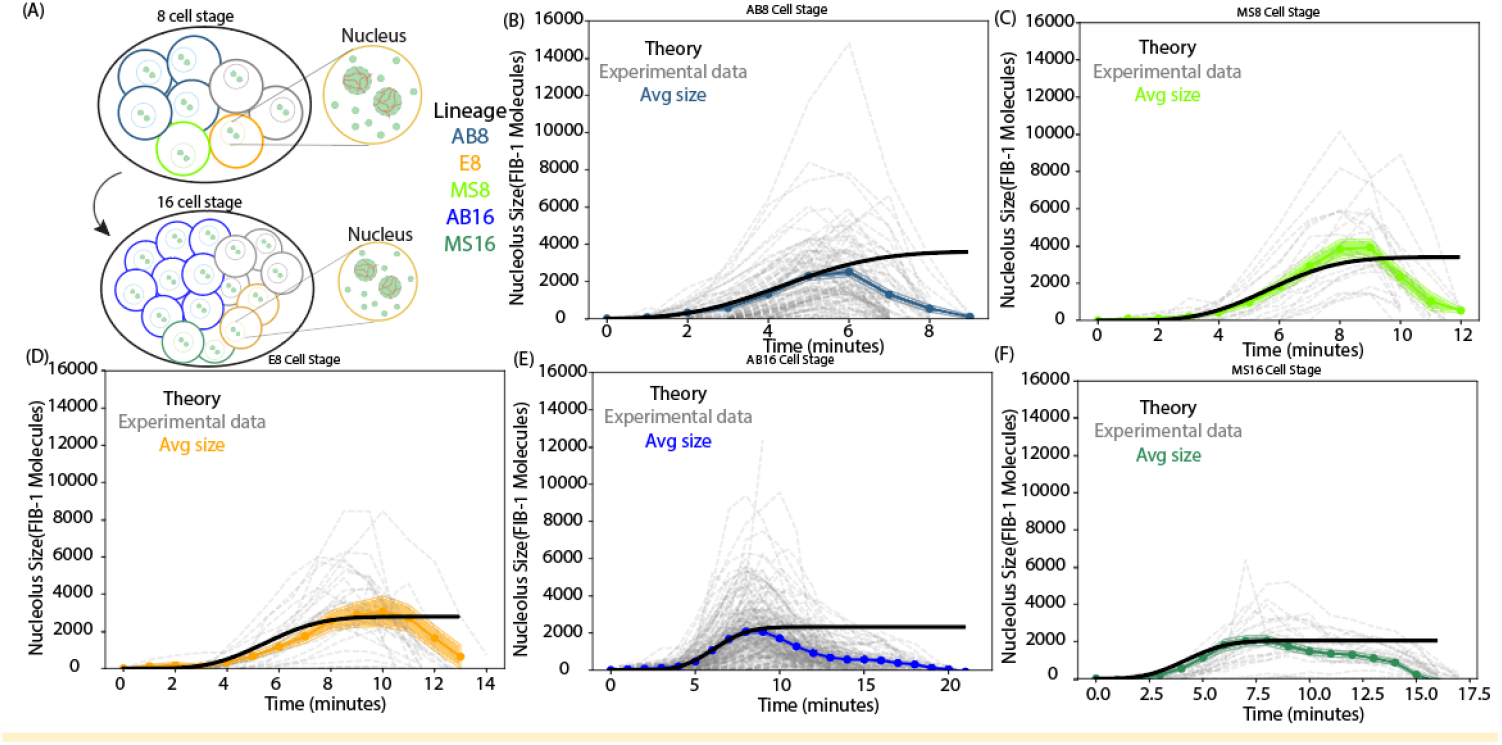
Nucleolar assembly at different stages of development. (A) Schematic for cell division from the 8 to 16-cell stage in *C. elegans* embryo. Two nucleoli assemble in each nucleus. Colors indicate different lineages: AB (blue), MS (green), E (orange). (B)-(F) Comparison of nucleolar size trajectories predicted for active transcription-templated assembly with data from fluorescence experiments. The experimental data show the raw trajectories (gray), the mean (dots) and standard error of the mean (shaded) over different independent measurements. Number of trajectories measured for AB8 = 87, MS8 = 22, E8 = 24, AB16 =180 and MS16 = 24.

When comparing the predictions of the active model to the data (Figures 3 B-F), we observe that for all the cell stages the nucleoli start disassembling shortly after the steady state size is reached. We also observe, that while the trajectory predicted by the active model (black line) matches the trajectories for the average size of the nucleolus for each lineage (colored lines), there is large variation in the individual nucleolar trajectories (gray-dotted lines) measured in experiments for all lineages. These observations are taken up again in the Discussion and the SI.

To study our model of active transcription-templated assembly in the case of cell-volume changes, we used RNAi to change embryo size, while keeping the number of nucleolar components fixed. Previously, it was shown that knockdown of the anillin homolog ANI-2 (15) or importin *α* IMA-3 (16) resulted in embryos that are smaller than control embryos (25% and 55%, respectively), while knockdown of the gene C27D9.1 (17) results in embryos 55% larger than control (7). As expected from the model, smaller (bigger) embryos had larger (smaller) nucleoli at maximum intensity (7). Here, we study the dynamics of the growth for nucleoli in the AB lineage for each RNAi condition at the 8-cell stage. Assuming that RNAi perturbations only change nuclear volume, we compute theoretical predictions for the nucleolar size *M*(*t*) for three RNAi conditions using parameters *N, ka* and *K*_*d*_ for AB8 lineage and corresponding nuclear volume. We find that the final size of the nucleoli decreases as the volume increases (Figure 4B, also reported in ref (7)) and also, the growth dynamics are faster for the cells with smaller nuclear volume. The increase in speed of assembly for cells with smaller volume is borne out by experiments (Figure 4C).

**Figure 4:**
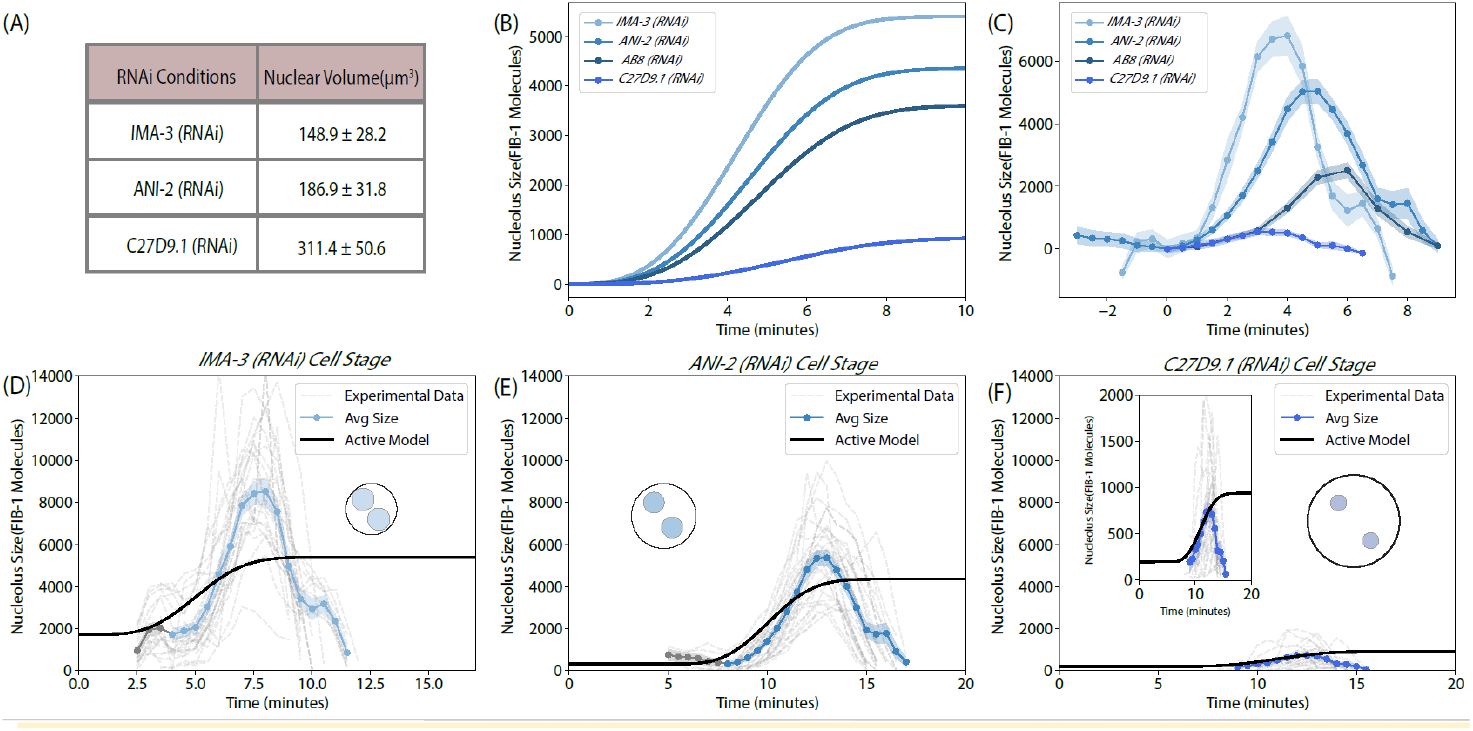
Comparing model predictions with RNAi conditions. (A) Measured nuclear volume for *ima-3(RNAi), ani-2(RNAi)* and *C27D9*.*1(RNAi)*. (B) Predicted nucleolar size trajectories from the active transcription-templated assembly model for AB8 cells in control and RNAi conditions. (C) Mean (points) and standard error of the mean (shaded) nucleolar size trajectories measured in experiments, with rescaled time. (D-F) Comparison of experimental data and theoretical prediction. Shaded regions are SEM errors. Number of trajectories measured for *ima-3(RNAi)* = 31, *ani-2(RNAi)* = 28 and *C27D9*.*1(RNAi)* = 24.

We find that the predicted nucleolar trajectories for different RNAi conditions (black lines in Figure 4 D-F) are consistent with measurements for two out of the three conditions (Figures 4D and F), and that all the nucleoli start disassembling around the time they reach their predicted steady state size. We discuss these results in the next section while analyzing the assumptions of the model and propose new experiments to further scrutinize the active mechanism of nucleolar assembly.

## Discussion

In early worm embryos, nucleolar proteins first condense into several droplets (ENDs), two of which become the nucleolus. Experiments also show that rRNA transcription is required for nucleolar assembly, whereas in the absence of transcription, only ENDs persist (6). Inspired by these experiments and the likelihood that in early embryos, nucleolar assembly happens on the same timescale as rRNA transcription, we considered two models of transcription-templated nucleolar assembly: one where the nucleolus assembles on previously transcribed rRNA, (Passive transcription-templated assembly) and another where the simultaneous process of rRNA transcription drives nucleolus assembly (Active transcription-templated assembly). While ENDs seem to form initially in the WT embryo (presumably before transcription of the rRNA start) and disappear by transferring their mass to the two nucleoli, our models describe the growth of the nucleoli within the nucleus, in order to make connection with experimental data.

We compare predictions of the two models of transcription-templated nucleolar assembly with fluorescence data measured in experiments. Remarkably, we find that while both models correctly predict the size of nucleoli at steady state, only the model of active assembly is able to capture the rapid kinetics observed in early embryos at different developmental stages, and for different RNAi perturbations of the embryo size. Our results highlight the importance of studying not only the properties of assembled organelles in steady state, but also the dynamics of assembly, in order to reveal the mechanisms of assembly.

### Scaling of nucleolar size with time

The active transcription-templated assembly model makes several quantitative predictions about the growth of the nucleolus in early worm embryos, which can be used to further test the model. From the solution of equation 5, we note that the characteristic time for growth is 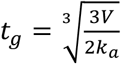. This parameter is about 5 min for the cell stages and lineages considered here, as nuclear volume *V* only varies from 100 − 200 *μm*^3^ and the estimated value of 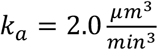. In the limit of *t* ≪ *t*_*g*_, using equation 5, we predict that 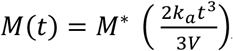, whereas for passive model, when 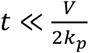 (about 5 min for the cell stages and lineages), 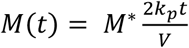, where 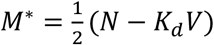 is identical for both models. These expressions further predict that if the nucleolar data is rescaled as 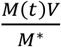 then we obtain a different scaling with time for the two models, given by

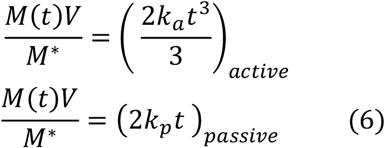

In Figure 5, we plot size ratio 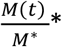 Nuclear Volume *V* as a function of time for each lineage (5A) and RNAi conditions (5B) and observe that the growth curves from different cell types collapse onto a single curve at short times, and seem to scale with time to the third power (See methods for details of data rescaling), as predicted by the active model.

**Figure 5:**
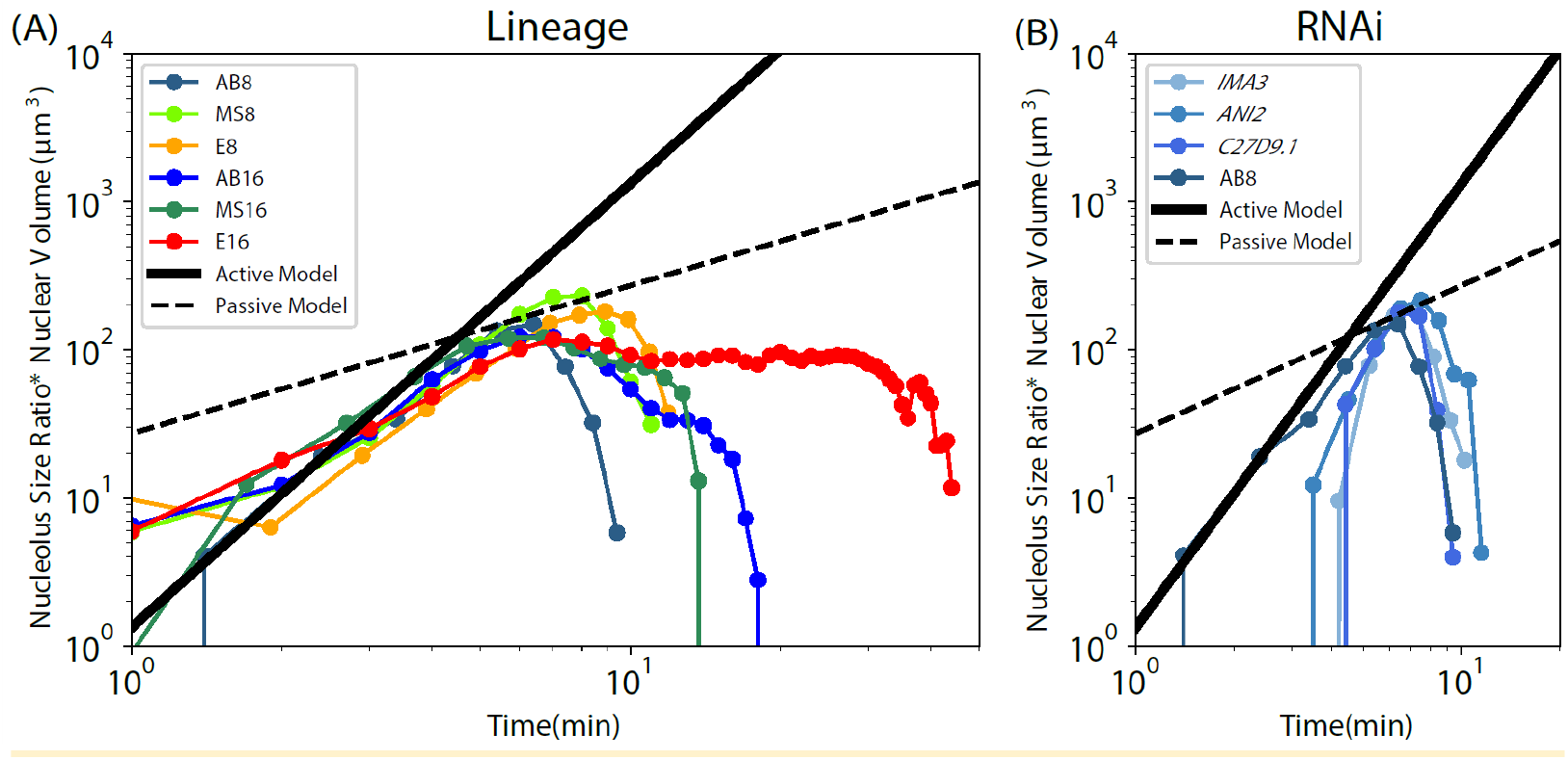
Growth scaling. Growth curves of all stages, (A) lineages and (B) RNAi conditions are rescaled to relative size 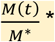 nucleus volume V. We compare with the theoretical prediction of scaling with time predicted in equation 6, by active (black solid line) and passive model (black dotted line).

Further, equation 6 predicts that the growth rate can be increased by reducing the parameter *t*_*g*_ - by either reducing the volume (like different RNAi conditions) or increasing transcription rates. Analogously, we predict that reducing transcription rates would slow the assembly which could be a way to further test the proposed model. A previous study had shown that knocking down the RNA polymerase I transcription initiation factor *C36E8*.*1*, which is essential in yeast (RRN3) (18) and mice (TIF-1A) (19), resulted in the nucleolus not forming (6). We propose that a systematic titration of this initiation factor would result in a decrease initiation rate and hence a smaller overall transcription rate (and hence smaller growth rate). Another way to reduce transcription would be to reduce the number of copies of rDNA units which are transcribed to rRNA, as has been done previously in yeast and bacteria (20, 21), and recently in *C. elegans* (22).

### Maximal size of nucleolus is close to the steady state size predicted by the model

Both transcription-templated assembly models described here predict that the steady state size of a nucleolus only depends on the number of molecules, their dissociation constant (or, equivalently, the critical concentration for condensation), and the volume of the cell. Notably, our description shows that the maximal nucleolar sizes measured in a previous work (7) (in AB8 cells) are close to the steady state size predicted by the models. Interestingly, from our theoretical analysis, we notice that even in cells from different lineages at the 8 and 16-cell stages, (with the exception of E16 cells) nucleoli reach their predicted steady state size and dissolve shortly after, presumably, in preparation for cell division. This observation raises intriguing questions about the synchrony of nucleolar assembly, cellular mechanisms driving dissolution of nucleolus, and nuclear envelope break down preceding cell division in these cells. Concurrently, we also note a large variation in nucleolar size from individual measurements which our analysis, describing the average behavior, cannot address. Whether this is due to population level noise arising from variation in nuclear sizes (Figure S5), or the stochastic nature of transcription remains an interesting open question.

### Deviations from model predictions for RNAi perturbations

In order to compare our model with data from RNAi perturbations, we assumed that only the nuclear volume is changed in our model and use parameters from control AB8 cells. While the predicted mean nucleolar trajectory agrees reasonably well for *ani-2(RNAi)* and *C27D9*.*1(RNAi)*, the *ima-3(RNAi)* data show interesting deviations (Figure 4 and 5B). First, we observe that nucleoli in *ima-3(RNAi)* grow even faster than predicted by the active model. We also note that for *ima-3(RNAi)* and *ani-2(RNAi)*, there is an overshoot in nucleolar size, where the measured maximum nucleolar intensity is higher than the steady-state size predicted by the model. We speculated whether this is due to the uncertainty in the measurement of nuclear volume or the estimation of the assembly parameter *k*_*a*_ (Figure S3).

Since our model relies on sample volumes measured for different conditions, we calculated a prediction band for possible nucleolar size, taking into account the maximum and minimum values of measured volume. Interestingly, we find that while the predictions for steady state are within the limits of experimental data for *ani-2(RNAi)* and *ima-3(RNAi)*, we predict negative nucleolar size for *C27D9*.*1(RNAi)*, corresponding to the maximum value measured for nuclear volume. This is most likely due to population level noise in the volume measurement which is larger than what is expected due to measurement error. Also, *C27D9*.*1(RNAi)* has the maximum deviation in the comparison between theory and experiments for the scaling of size ratio 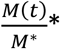 Nuclear Volume *V* (Figure 5B). Since this rescaling relies on the measurement for initial size and delay time *t*_0_ (See Methods), we believe that this discrepancy could be attributed to the small size of nucleoli in this condition (below the diffraction limit), which most likely are not being detected in the early stages of growth.

We note that the fast growth in *ima-3(RNAi)* remains an interesting deviation from the active model prediction and cannot be explained by uncertainty in the measurement of nuclear volume or the estimation of the assembly parameter *k*_*a*_ (Figure S4). IMA-3 is a nuclear import factor (16), and so it is possible that this knockdown alters the composition of the nucleoplasm by disrupting nucleocytoplasmic transport, which could affect rRNA transcription in an indirect way. Additionally, knockdown of *ima-3* is also embryonic lethal, while *ani-2* and *C27D9*.*1* RNAi embryos hatch and develop into viable, fertile worms. It is possible that the nuclear volume in this condition may be small enough to cause other volume effects that are not considered in the model. Tracking the production of rRNA for control and *ima-3(RNAi)* embryos will be crucial in examining this hypothesis.

### RNA-mediated growth of condensates

Transcription can affect condensate assembly in a variety of ways, as has been explored in many previous studies (11, 23–25). These works fall under two categories – one that relies on a thermodynamic equilibrium between a protein when it is bound and unbound to RNA, and the other where the process of transcription itself is actively influencing the assembly of condensates. An example of mRNA-induced assembly is the two distinct types of Whi3 droplets seen in *Ashbya* cells (13), where those clustered around nuclei and the growing tips have different amounts of Whi3 incorporation. Distinct Whi3 binding mRNAs segregate to different droplets in a common cytoplasm. Another example is the *in vitro* assembly of FIB-1 condensates reported in (6), in which addition of RNA both lowers the saturation concentration and accelerates droplet growth. Importantly, these FIB-1/RNA droplets behave similarly to ENDs – but not nucleoli – in terms of their assembly kinetics; they do not exhibit the concavity at early times, which is characteristic of nucleoli (Fig. 2B, 3B-F, 4C). All of these examples rely on the rate of assembly being accelerated by the presence of RNA. In contrast and similar to the nucleolus, nascent RNAs have been shown to stimulate condensate formation while bursts of RNAs produced during elongation accelerate the dissolution of condensates (11). Further, methylation of nascent RNAs has been shown to facilitate the formation of transcriptional condensates (12).

Inspired by these experiments, many theoretical studies have explored the interaction between transcription and condensate assembly using mathematical modeling. Recently, Joseph et al used a patchy-particle polymer model to investigate liquid-liquid phase separation of protein-RNA mixtures and found that RNA can indeed accelerate the nucleation stage of phase separation (24). In contrast, Schede et al described how gene activity can precisely localize and nucleate nuclear condensates (25). Their theoretical model, which described how the processes of RNA synthesis, degradation, and diffusion shape the dynamics and organization of nuclear condensates, which are themselves composed of RNAs. This work explores an interesting feedback where the transcribing gene codes for a transcription factor for the condensate protein itself, and studies its effect at steady state. Our work, in contrast, describes a situation where the transcribing gene is responsible for creating more nucleating sites for the nucleolus to grow, and thereby affecting the kinetics of its assembly. In that way, it can be described as an initial value problem in development - where the process of transcription can not only “time” but in some cases, especially where it is nascent, also “localize” the process. A quantitative study of each of these processes will enable us to analyze the diversity of ways in which transcription can influence the assembly of membraneless organelles in cells.

In this paper, we described a model of transcription-templated assembly of the nucleolus. We used previously published experimental results to study the role of transcription of rRNAs in the assembly of the nucleolus in early *C. elegans* embryos at different cell stages. We demonstrate that our model of active transcription-templated assembly quantitatively describes the relationship between nascent rRNAs and size of nucleolus. Excitingly, it predicts the early kinetics of assembly seen in experiments, and that nucleolar-size data from cells at different stages of development can be collapsed onto a single master curve. It also makes testable predictions for future experiments, which relate transcription dynamics, in particular the rates of initiation and elongation of rRNA, to the dynamics of nucleolar assembly.

Our work is an example of how quantitative models can help us understand the role of different mechanisms in the growth process during development. In this work, we focused on the assembly kinetics and did not consider mechanisms influencing the dissolution of the nucleolus which can be cell cycle dependent – mediated by the cellular cyclin-dependent kinase (26) or depend on the local concentration of nucleolar proteins which can change due to nuclear import (7) and/or nuclear volume changes (6). A more detailed model accounting for each of these observations will enable us to understand the full assembly and disassembly of the nucleolus over a complete cell cycle.

## Acknowledgements

We would like to thank members of Mohapatra, Kondev and Weber groups for stimulating discussions. This work was supported by the National Institute Of General Medical Sciences of the National Institutes of Health R35GM147556 (L.M. and N.K.), the National Science Foundation grants DMR-1610737 and MRSEC DMR-2011846, and by the Simons Foundation (J.K.), and Canadian Institutes of Health Research (PJT159580) to S.C.W.

