## Supplementary material for "Transcription templated assembly of the nucleolus in the *C. elegans* embryo": Supp

**Supplemental Information/Methods:**

### 1. Estimation of number of molecules (N) and dissociation constant ($K_d$) in AB lineage at 8-cell stage

To estimate model parameters $N$ and $K_d$, we focused on Figure 2B in ref(7) which reported the total fluorescence of FIB-1 proteins in both nucleoli of AB-lineage cells at the 8-cell stage for different RNAi conditions. We first used the fluorescence to number of molecules conversion parameter called α in the SI(7) and converted the intensity into number of FIB-1 molecules. Our models predict the total nucleolar size at steady state $M^*_{tot} = (N - K_d V)$, where $V$ is the volume of the nucleus and $N$ is the number of FIB-1 molecules in the nucleus. We used Matlab's fitting function and Maximum Likelihood Estimation (Figure 2A and S1) to obtain parameters for AB8 cells: $N^{AB}_{8-cell} = (1.9 \pm 0.1) \times 10^4$ molecules, and $K_d = 55 \pm 6\ mol/\mu m^3$, values which are in the same range as reported in (7). By assuming that the parameter $K_d$ and the nuclear concentration of FIB-1 remain the same for all lineages at 8 and 16-cell stages, we are able to predict the total number of molecules $N$ pertaining to all cell types (See Table 1).

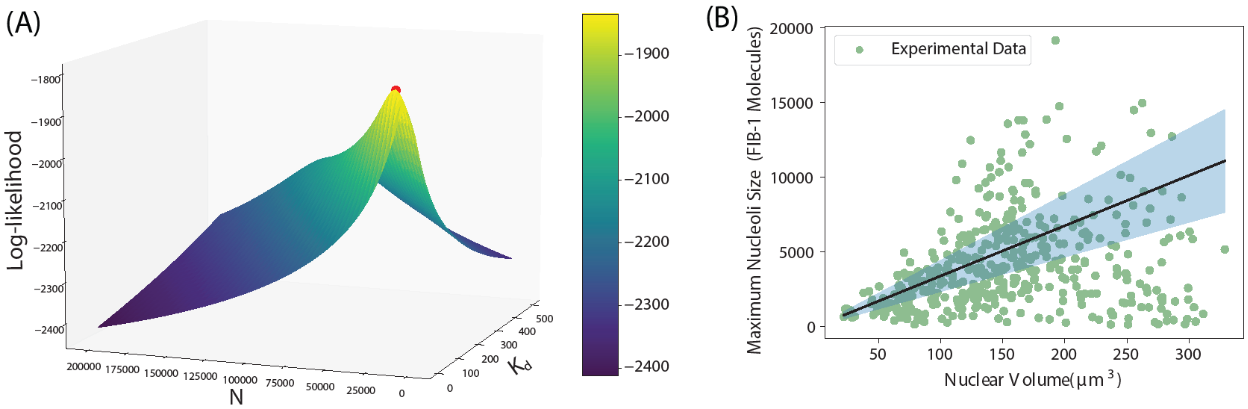


**Figure S1: Extracting parameters N and $K_d$ from published data for AB8 cells** (A) 3D projection of the Log-Likelihood function is plotted against $N$ and $K_d$ values, using published data (ref 7) from AB8 cells. The peak (in red) corresponds to best parameter fits $N^{AB}_{8-cell} = (1.9 \pm 0.1) \times 10^4$ molecules, and $K_d = 55 \pm 6\ \mu m^{-3}$. (B) Published data for nucleolar size of AB cells across development is fitted against direct scaling equation $(C - K_d)V$, where $C$= saturation constant, and $V$ = nuclear volume. We obtain $C$, and use it to estimate errors in estimation of $N$ for other lineages.

### 2. Errors in the estimation of N for all lineages:

For 8-cell stage, using published data (Figures 2A and S1) we estimate $N_{AB} \approx 19000 \pm 1463$. The FIB-1 protein concentration C for the AB8 cell lineage is calculated using the ratio of the number of protein molecules N to the nuclear volume V. The value for NAB8 was obtained via maximum likelihood estimation, while VAB8 was derived from experimental measurements (Table 1). Using error propagation, $\frac{\Delta C}{C} = \sqrt{\left(\frac{\Delta N}{N}\right)^2 + \left(\frac{\Delta V}{V}\right)^2}$ . Using $\Delta N = 1463$, $\Delta V = 36.4\ \mu m^3, N = 19000$ and $V = 214.3\ \mu m^3$, $C_{AB8} = 88.66 \pm 16.53\ molecules\ \mu m^{-3}$. We assume that the concentration of FIB-1 proteins remains constant within the embryo, and use measured volumes in Table 1 to calculate the number of molecules and their associated error. For each lineage i, number of molecules $N_i = C \times V_i$ and $\frac{\Delta N_i}{N_i} = \sqrt{\left(\frac{\Delta C}{C}\right)^2 + \left(\frac{\Delta V_i}{V_i}\right)^2}$. These are reported in Table 1.

### 3. Estimation of assembly parameters for Passive and Active transcription-templated assembly using E16 data

Next, we use the early assembly portion of the nucleolar size trajectories from E lineage cells at 16-cell stage to estimate parameters for both models. Unlike the other lineages and previous stage, in which nucleoli dissolve after reaching a maximum size, these nucleoli appear to hover around the same size for ~ 20 min before dissolving prior to cell division (Figure 2D). Using $N$ and $V$ for E16 cells, we obtain $k_p = 13.6 \pm 1.8\ \mu m^3 min^{-1}$ for the passive model and $k_a = 2.0 + 0.5 \frac{\mu m}{min^3}$ for the active model (Figure S2). Comparing the predicted curves, we note that the two models show different time dependence while leading to the same steady state size. Remarkably, the active model is able to capture the concavity seen in early time, indicating indeed the rRNA template is not pre-formed but accumulates concurrently with the nucleolus.

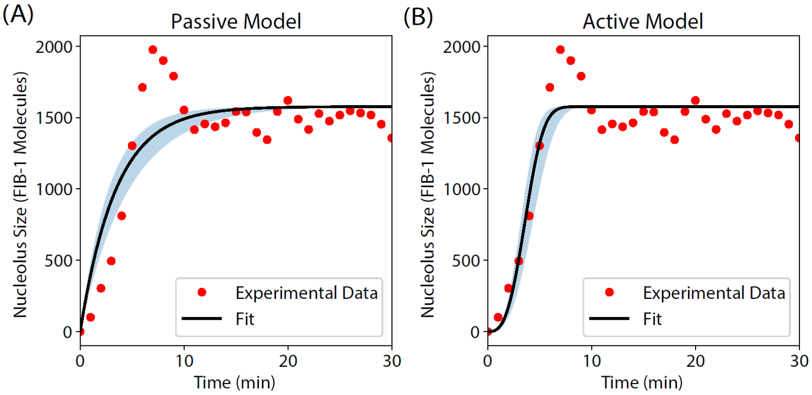


**Figure S2: Fitting theoretical models with experimental data from E16 cells.** Fits of the (A) Passive and (B) Active models (red lines) to experimental data (black points) from E16 cells. We use the Equation S5 (solution to Equation 3 and 4 in main text) for the fitting, and obtain $k_p = 13.6 \pm 1.8\ \mu m^3 min^{-1}$ for the passive model and $k_a = 2.0 \pm 0.5 \frac{\mu m}{min^3}$ for active model.

### 4. Comparing with other lineages in worm embryo

Equation 2 in the main text describes the growth of the nucleolus for the model of Active transcription-templated assembly**,** where the nucleolus starts from zero initial time and size. In this section, we want to compare the analytical solution to experimental data obtained from fluorescence experiments for all lineages at the 8 and 16-cell stages. All the data sets had different initial size and time. All times were measured relative to nuclear envelope breakdown (NEBD) in AB-lineage cells at the 4-cell stage, so we first rescaled time in experimental data by the first data time point. Since the first data point may not be the time zero for nucleolus, we solved equation 2 using initial conditions where each nucleolus is starting from an initial size $M_0$ and time $t_0$. The time-dependent solution for nucleolar size $M(t)$ to the differential equation in that case is given by

$$M(t) = M_0 e^{-\frac{2k_a(t-t_0)^3}{3V}} + \frac{(N-k_dV)}{2}\left(1 - e^{-\frac{2k_a(t-t_0)^3}{3V}}\right) \quad (S1)\ .$$

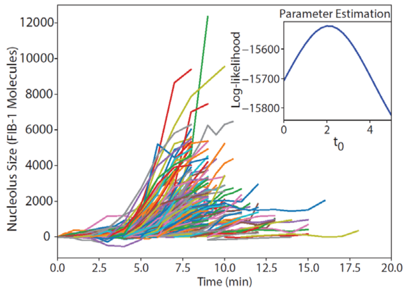


**Figure S3: Maximum Likelihood Estimation for $t_0$.** We parse the early-time part of nucleolar trajectories (AB 16 shown here) and use log-likelihood to estimate $t_0$ values for all lineages, stages and RNAi conditions.

| Label | $t_0$ | Number of Trajectories |
|---|---|---|
| AB8 | -0.4 ± 0.1 | 87 |
| MS8 | 1.0 ± 0.2 | 22 |
| E8 | 1.1 ± 0.2 | 24 |
| AB16 | 2.1 ± 0.1 | 180 |
| MS16 | 1.3 ± 0.3 | 24 |
| *IMA-3(RNAi)* | -3.2 ± 0.2 | 29 |
| *ANI-2(RNAi)* | -2.5 ± 0.1 | 28 |
| *C27D9.1(RNAi)* | -3.4 ± 0.2 | 23 |

**Table S1:** Estimated value of transcription start time for cell types and RNAi conditions, along with the number of nucleolar trajectories.

We fit this equation to experimental data of early assembly to obtain a lineage-specific parameter $t_0$, which corresponds to the time difference between "time zero" of nucleolar assembly (i.e. start of transcription) and the first measured data point. We used a Maximum Likelihood estimation (Figure S3) to obtain $t_0$ (measured relative to the first data point) and find that, remarkably, the range of time $t_0$ for most WT lineages is within $\pm$ 1 min (Table S1), which is the frequency of measurement. This suggests that the first experimental measurement is very close to the emergence of nucleoli in all lineages considered here. The comparison of model prediction to experimental results is plotted in Figures 3 (with SEM) and S4 (SD).

When $t < \sqrt[3]{\frac{3V}{2k_a}}$, Equation S1 can be written as

$$M(t) - M_0 = \left(\frac{(N - K_dV)}{2} - M_0\right)\left(\frac{2k_a(t - t_0)^3}{3V}\right) \quad (S2)$$

where $M_0$ is the first measured nucleolar size.

Comparing this to the passive model, the time-dependent solution for nucleolar size $M(t)$ where each nucleolus is starting from an initial size $M_0$ and time $t_0$, is then given by

$$M(t) = M_0 e^{-\frac{2k_p(t-t_0)}{V}} + \frac{(N-k_dV)}{2}\left(1 - e^{-\frac{2k_p(t-t_0)}{V}}\right) \quad (S3)\ .$$

At $\ll \frac{V}{2k_p}$, we can approximate $e^{-\frac{2k_p(t-t_0)}{V}} = 1 - \frac{2k_p(t-t_0)}{V}$ and hence

$$M(t) = M_0\left(1 - \frac{2k_p(t-t_0)}{V}\right) + \frac{(N - k_dV)}{2}\left(\frac{2k_p(t-t_0)}{V}\right)$$

$$M(t) - M_0 = \left(\frac{(N - K_dV)}{2} - M_0\right)\left(\frac{2k_p(t-t_0)}{V}\right) \quad (S4)$$

Note that if $M_0$ and $t_0$ are ignored, then we reproduce the results for the passive and active models shown in the main text, i.e.

$$M_{passive}(t) = \left(\frac{N - K_dV}{2}\right)\left(1 - e^{-\frac{2k_pt}{V}}\right)$$

$$M_{active}(t) = \left(\frac{N - K_dV}{2}\right)\left(1 - e^{-\frac{2k_at^3}{3V}}\right) \quad (S5)$$

All the data was rescaled $M \rightarrow M - M_0$ and $t \rightarrow t - t_0$ to compare with the active model prediction in Figure 3. The comparison of data with SD is shown in Figure S4. Nuclear volumes were measured in bulk and had a SD associated with them. In Figure S5, we study the effect of uncertainty in our estimation of $k_a$ and nuclear volume measurement on the predicted nucleolar size.

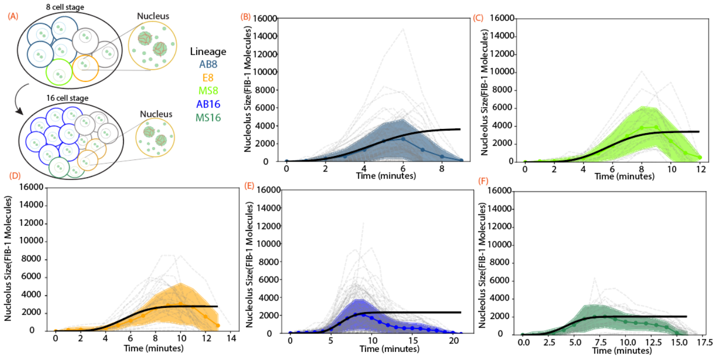


**Figure S4: Nucleolar assembly at different stages of development** (B)-(F) Comparison of simulated nucleolus size trajectories with data from fluorescence experiments for active transcription-templated assembly. Raw trajectories in dotted – gray, and mean shown in solid colors with SD as shaded regions. Number of trajectories recorded for AB8 = 87, MS8 = 22, E8 = 24, AB16 =180, and MS16 = 24.

Equation S2 and S4 further predict that $\left(\frac{M(t)-M_0}{M^*-M_0}\right)V$, where $M^* = \frac{1}{2}(N - K_dV)$, has a distinct dependence on time, for the two models. For the active model $= \frac{2k_a(t-t_0)^3}{3}$, whereas it is $2k_p(t - t_0)$ for the passive model. These are the expressions used in Figure 5.

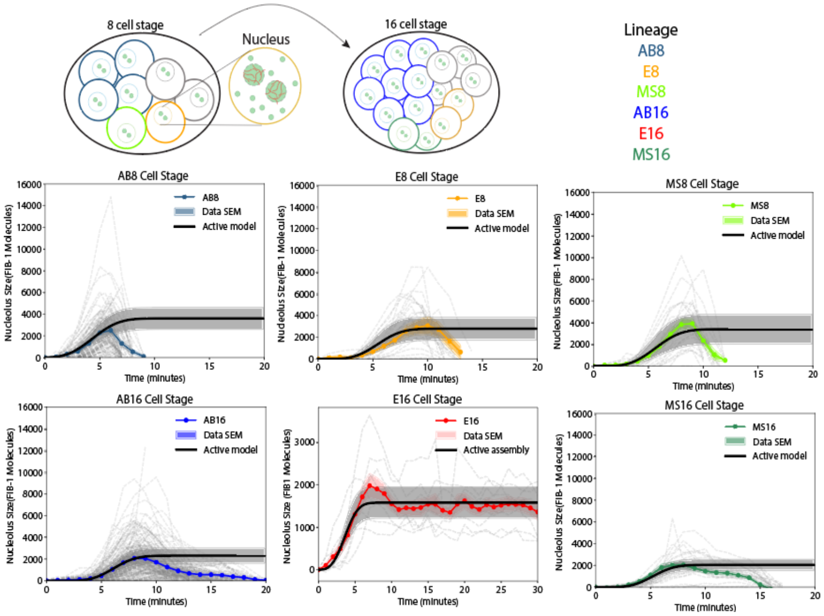


**<u>Figure S5: Effect of uncertainty in nuclear volume</u>** (A)-(F) Active Model Comparison (black) with experimental data (error bars represent SD) for different lineages while accounting for uncertainty in estimation of nuclear volume. Shaded regions correspond to uncertainty in size estimation. Variation in nuclear volumes is listed in Figure 2A. Raw trajectories in dotted – gray, and mean shown in solid colors. Number of trajectories recorded for AB8 = 87, MS8 = 22, E8 = 24, AB16 =180, and MS16 = 24.

### 5. Comparison with RNAi Data

We performed an MLE analysis RNAi conditions to calculate $t_0$ for all RNAi conditions (Table S1) The larger deviation in $t_0 = 3.4\,min$ in *C27D9.1(RNAi)* is likely due to its small nucleoli being below the diffraction limit, and not being detected in early stages. This experimental limitation affects our estimate of initial size $M_0$ and hence the time of transcription initiation, $t_0$.

In Figure S6, we study the effect of uncertainty in our estimation of $k_a$ and nuclear volume measurement on the predicted nucleolar size.

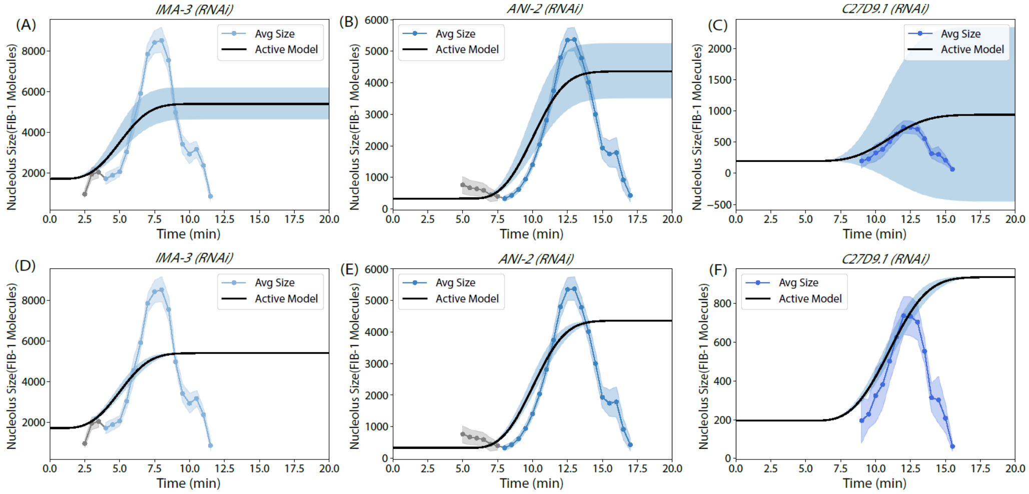


**<u>Figure S6: Effect of uncertainty in $k_a$ and nuclear volume</u>** Active Model Comparison (black) with experimental data (error bars represent SD) from RNAi conditions while accounting for uncertainty in estimation of $k_a$ (A)-(C) and variation in nuclear volume measurement (D)-(F). Shaded regions correspond to uncertainty in size estimation. Uncertainty in $k_a$ is represented by $2.0 \pm 0.7 \frac{\mu m}{min^3}$, and while variation nuclear volumes is listed in Figure 4A. Number of trajectories considered for *ima-3(RNAi)* = 31, *ani-2(RNAi)* = 28 and *C27D9.1(RNAi)* = 24.

### 6. Estimate of Timescales of rRNA transcription

Using the length of an rDNA gene unit = 10 kb, and the rate of transcription = 1 kb/min, we estimate the time to reach the end ∽ 10 min, which is similar to nucleolar assembly times.

### 7. Acquisition of time-lapse movies of nucleolar assembly in early embryos

This workflow was used in an earlier study (7). Nucleoli were visualized using a transgenic worm line expressing FIB-1::GFP from a fosmid that was randomly integrated into the genome (Clone 445429768005125 D03 (27). Worms were maintained at 20oC on NGM plates seeded with OP50 or HT115(DE3) bacteria, for the lineage and RNAi datasets, respectively. Embryos were dissected from gravid hermaphrodites and mounted on M9-1% agarose pads. Timelapse movies were acquired on a two-photon laser scanning system custom-built around an upright Olympus BX51 microscope with an excitation wavelength of 960 nm. Emitted light was collected with a 40X/NA0.8 water immersion objective and an NA1.3 oil immersion condenser, and detected with high quantum efficiency GaAsP photomultiplier tubes (Hammamatsu). 3D volumes were acquired using an objective piezo controlled by ScanImage software (28).

RNAi experiments were performed by picking L4 larvae onto NGM plates containing 1 mM IPTG and 100 μg/mL ampicillin, seeded with feeding clones from the Ahringer library (29). The empty vector L4440 served as a negative control for all experiments. Worms were allowed to feed at 20°C for 36-48 hr before embryos were harvested for imaging.

Nucleoli were segmented from raw z-stacks using a 3D bandpass filter and objects were linked into trajectories using the Matlab Particle Tracking Code Repository (https://site.physics.georgetown.edu/matlab/index.html). Nucleolar size was calculated from the summed fluorescence intensity within each segmented object, which was converted to number of molecules by calibrating pixel intensity with a solution of known concentration of purified GFP, such that a = $3.1x10^3$ intensity units/pixel/uM.
